# Rcs phosphorelay activation in cardiolipin-deficient *Escherichia coli* reduces biofilm formation

**DOI:** 10.1101/522219

**Authors:** Julia F. Nepper, Yin C. Lin, Douglas B. Weibel

## Abstract

Biofilm formation is a complex process that requires a number of transcriptional, proteomic, and physiological changes to enable bacterial survival. The lipid membrane presents a barrier to communication between the machinery within bacteria and the physical and chemical features of their extracellular environment, and yet little is known about how the membrane influences biofilm development. We found that depleting the anionic phospholipid cardiolipin reduces biofilm formation in *Escherichia coli* cells by as much as 50%. The absence of cardiolipin activates the Rcs envelope stress response, which represses production of flagella, disrupts initial biofilm attachment, and reduces biofilm growth. We demonstrate that a reduction in the concentration of cardiolipin impairs translocation of proteins across the inner membrane, which we hypothesize activates the Rcs pathway through the outer membrane lipoprotein RcsF. Our study demonstrates a molecular connection between the composition of membrane phospholipids and biofilm formation in *E. coli* and suggests that altering lipid biosynthesis may be a viable approach for altering biofilm formation and possibly other multicellular phenotypes related to bacterial adaptation and survival.

**Importance:** There is a growing interest in the role of lipid membrane composition in the physiology and adaptation of bacteria. We demonstrate that a reduction in the anionic phospholipid cardiolipin impairs biofilm formation in *Escherichia coli* cells. Depleting cardiolipin reduced protein translocation across the inner membrane and activated the Rcs envelope stress response. Consequently, cardiolipin depletion produced cells lacking assembled flagella, which impacted their ability to attach to surfaces and seed the earliest stage in biofilm formation. This study provides empirical evidence for the role of anionic phospholipid homeostasis in protein translocation and its effect on biofilm development, and highlights modulation of the membrane composition as a potential method of altering bacterial phenotypes related to adaptation and survival.

## Introduction

Bacteria are often found in multicellular communities referred to as biofilms (1, 2). Biofilms are ubiquitous and persistent structures with a complexity and economic impact that has drawn broad attention to studying (and disrupting) the processes underlying their development (3-5). Attachment of cells to a substrate is the initial step in biofilm formation, and is facilitated by extracellular organelles, including pili and flagella (1, 6-8). As the biofilm grows and matures, cells within the community produce and secrete a polymer matrix (i.e., extracellular polymeric substance, EPS; or extracellular matrix, ECM) that encases the microbial consortium, providing a three-dimensional structure and protecting the cells within from hostile and fluctuating environments (e.g., dessication, shear forces, and the presence of antibiotics, antiseptics, and oxidants) (9, 10). ECM is dispensable for the first steps of biofilm formation in several bacteria (e.g., *Escherichia coli*) and required for later stages of development (11).

Many of the physiological changes that occur during biofilm formation involve processes associated with the cell membrane (e.g., an increase in secretion of polysaccharides or nucleic acids) or structures attached to it (e.g., pili and flagella). The cell membrane is a largely overlooked connection to biofilm-relevant factors and phenotypes. The membrane is a central participant in the activation of stress responses, including those propagated by the σ^S^ and Rcs (<u>r</u>egulation of <u>c</u>olanic acid <u>s</u>ynthesis)signaling pathways, which play important roles at different stages of biofilm development (12, 13). Many of the signaling pathways that impact biofilm development—e.g. Rcs and Cpx—involve proteins directly associated with the membrane and/or are activated by disruption of the lipid membrane (e.g. lipopolysaccharide defects) (14, 15). In addition, proteins that assemble into extracellular organelles are transported across and/or inserted into the cell envelope, participate in the attachment of cells to surfaces, and contribute to the ECM to enable biofilm formation. Consequently, alterations in the composition of bacterial membranes may impact the transport of these families of proteins.

A number of observations highlight the importance of bacterial membranes in biofilm formation, and recent work has deepened our understanding of the interplay of these two factors. Benamara *et al.* showed that the Gram negative opportunistic pathogen *Pseudomonas aeruginosa* undergoes significant changes in inner and outer membrane lipid composition when adapting to a biofilm state of growth, and other groups have elucidated important roles for rhamnolipid production at every stage of biofilm development in *P. aeruginosa* and *Agrobacterium tumefaciens* (16-19). Specific lipids present in the ECM are important in late stages of biofilm formation in *Mycobacterium tuberculosis* (20). In *Listeria monocytogenes*, biofilm attachment has been correlated with production of the fatty acids hexadecanoic acid and octadecanoic acid, which may be associated with or inserted into membranes (21). The connection between the composition and properties of membranes and biofilm formation is largely unstudied in the vast majority of bacteria, including the model gram-negative bacterium *Escherichia coli* (22).

*E. coli* cells growing exponentially under typical laboratory conditions and at exponential phase have cell membranes consisting of the following families of lipids and their relative concentration: 70% phosphatidylethanolamine (PE), 15-25% phosphatidylglycerol (PG), and 5-10% cardiolipin (CL) (Fig. 1) (23). The lipid composition of cell membranes changes as cells enter the stationary phase—e.g., the concentration of CL increases to 15-20% of total lipid (23)—however nothing is known about the changes that occur directly before or during biofilm formation.

**Figure 1.**
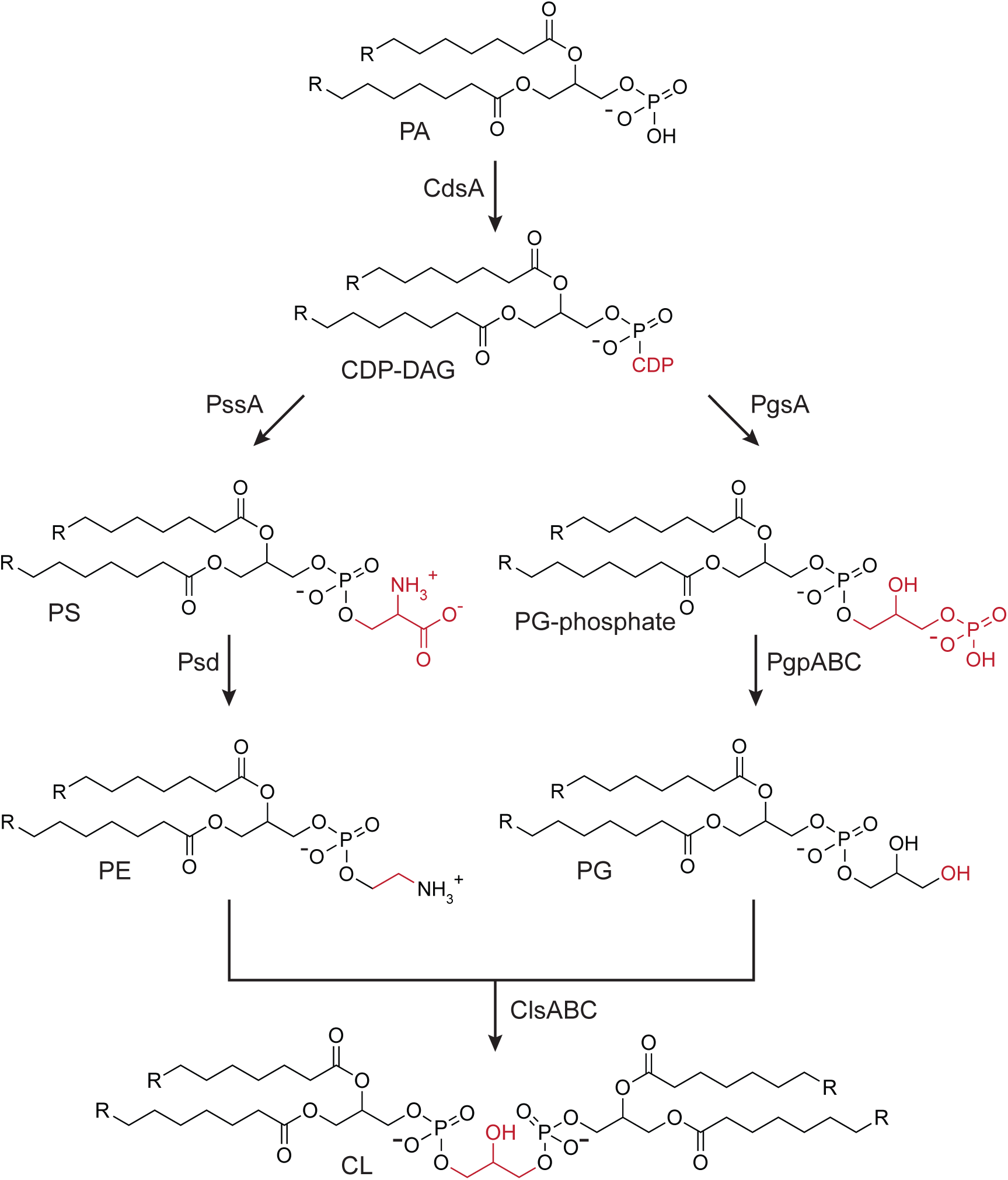
Phospholipid biosynthesis pathways in *E*. *coli*. Chemical transformations for CL biosynthesis in *E. coli* cells. Abbreviations for phospholipids are: PA, phosphatidic acid; CDP-DAG, CDP diacylglycerol; PS, phosphatidylserine; PE, phosphatidylethanolamine; PG, phosphatidylglycerol; and CL, cardiolipin. The lipid composition of wild-type *E. coli* growing exponentially in typical laboratory growth conditions (37°C in rich media with agitation) is 70% PE, 15-25% PG, and 5-10% CL.

CL is an unusual four-tailed anionic phospholipid synthesized by three nonessential phospholipase D type enzymes: <u>c</u>ardiolipin <u>s</u>ynthase A (ClsA), ClsB, and ClsC (23). The function of CL in bacterial physiology is not entirely known, however growing evidence supports a role for CL in localizing and activating proteins in cells (24-27). Recently, Rowlett *et al.* (2017) demonstrated cardiolipin synthases influence the attachment of *E. coli* cells to surfaces (22). Concomitant with that study, we discovered that depleting CL causes a drastic decrease in both early biofilm (here we refer to it as “surface attachment”) and late stage biofilm formation. In characterizing this phenotype, we discovered that disrupting CL biosynthesis activates the Rcs envelope stress response and leads to downstream phenotypes that alter biofilm formation.

Initiation of the Rcs signaling system leads to the phosphorylation of RcsB, enabling it to dimerize and function as a transcriptional regulator (28). Phosphorlyated RcsB can also form a heterodimer with the auxiliary protein RcsA (29); the RcsB-RcsB and RcsA-RcsB complexes control the expression of a number of genes involved in acid resistance, as well as the colanic acid and curli synthesis operons, osmotically inducible peroxidase *osmC*, and the small regulatory RNA rprA (30-32). We found that activation of the Rcs pathway has little or no effect on Δ*cls* cells growing planktonically.

Disrupting the Rcs system is sufficient to restore surface attachment in *cls* mutants, suggesting that Rcs activation is responsible for the biofilm defects we observe. The data we describe supports a model in which CL reduction impairs the translocation of proteins across the inner membrane (IM), which initiates the Rcs stress response, leading to a downstream reduction in biofilm formation.

## Results

### Cardiolipin affects surface attachment

Previous studies have established that in the exponential phase of growth, CL represents ∼5% of total lipid composition in *E. coli* cells. Under certain environmental conditions, such as high osmolarity, low pH, or upon entry into stationary phase, CL content increases by as much as 200% (33). As nothing was known at the onset of this study regarding how lipid composition changes in biofilm-associated cells, we extracted and quantified total lipids from cells grown statically in 96-well microplates. As expected, TLC analysis of cells grown in minimal medium revealed a significant increase in CL in stationary phase cells (17.6% of total phospholipids) compared to exponentially growing cells (6.6% of total phospholipids; Fig. 2). Similar to stationary phase cells, CL was enriched in biofilm-associated cells (19.2% of total phospholipids). Interestingly, the proportion of PE decreased to approximately half of the total phospholipid in both stationary phase and biofilm-associated cells and was accompanied by a concomitant rise in PG levels.

**Figure 2.**
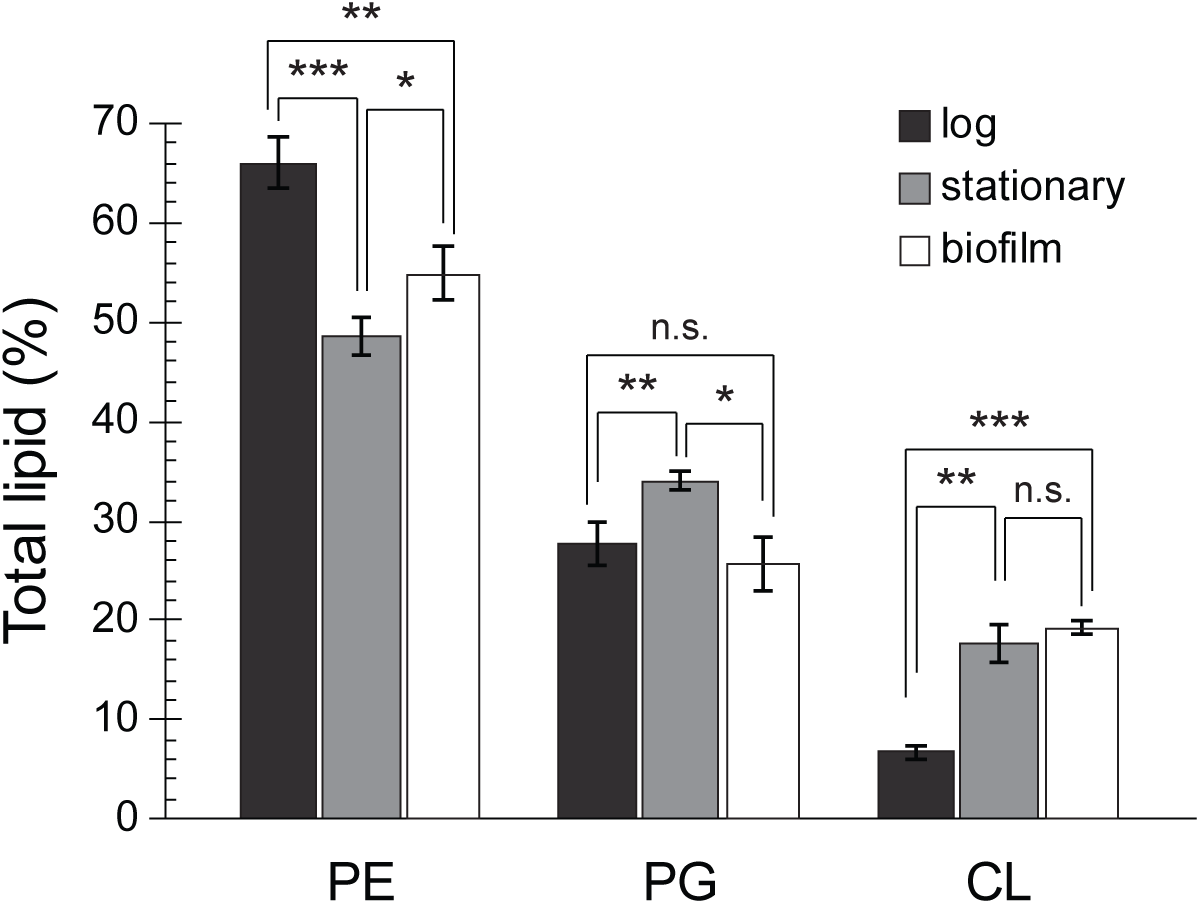
CL is elevated in cells from early stage biofilms compared to cells in log phase. Total membrane lipids were isolated from MG1655 cells grown in suspension to an absorbance (λ=600 nm) of 0.4-0.6 (log), for 24 h (stationary) or in 96-well plates without shaking for 24 h (biofilm). After separation by TLC, phospholipids were visualized by fluorescence imaging following treatment with cupric sulfate. ImageJ was used to quantitate lipid spots. Error bars indicate standard deviation of 3 biological replicates; \*\*\**p* < 0.001, \*\**p* < 0.01, \**p* < 0.05, n.s. not significant; Student’s *t* test.

To further investigate the effect of CL on biofilm formation, we used crystal violet to quantify adherence of cells lacking one or more *cls* genes to the surface of polystyrene microplates. In minimal medium—but not in LB—disrupting any combination of *cls* genes resulted in a significant (*p* < 0.05, unpaired *t* test) decrease in surface attachment relative to wild-type (WT) cells (Fig. 3*A*, Fig. S1). Even after incubation for 5 days, Δ*cls* cells produced less biofilm than WT cells (Fig S2). All three Cls enzymes (ClsA, B, and C) contain 2 phospholipase D-type HKD motifs, and mutation of one or both of these motifs renders Cls catalytically inactive. Using an arabinose-inducible expression vector to complement single Δ*cls* mutant strains, we were able to partially restore surface attachment by ectopic expression of WT, but not mutated *cls* (Fig. 3*B*, Fig *S*3).

**Figure 3.**
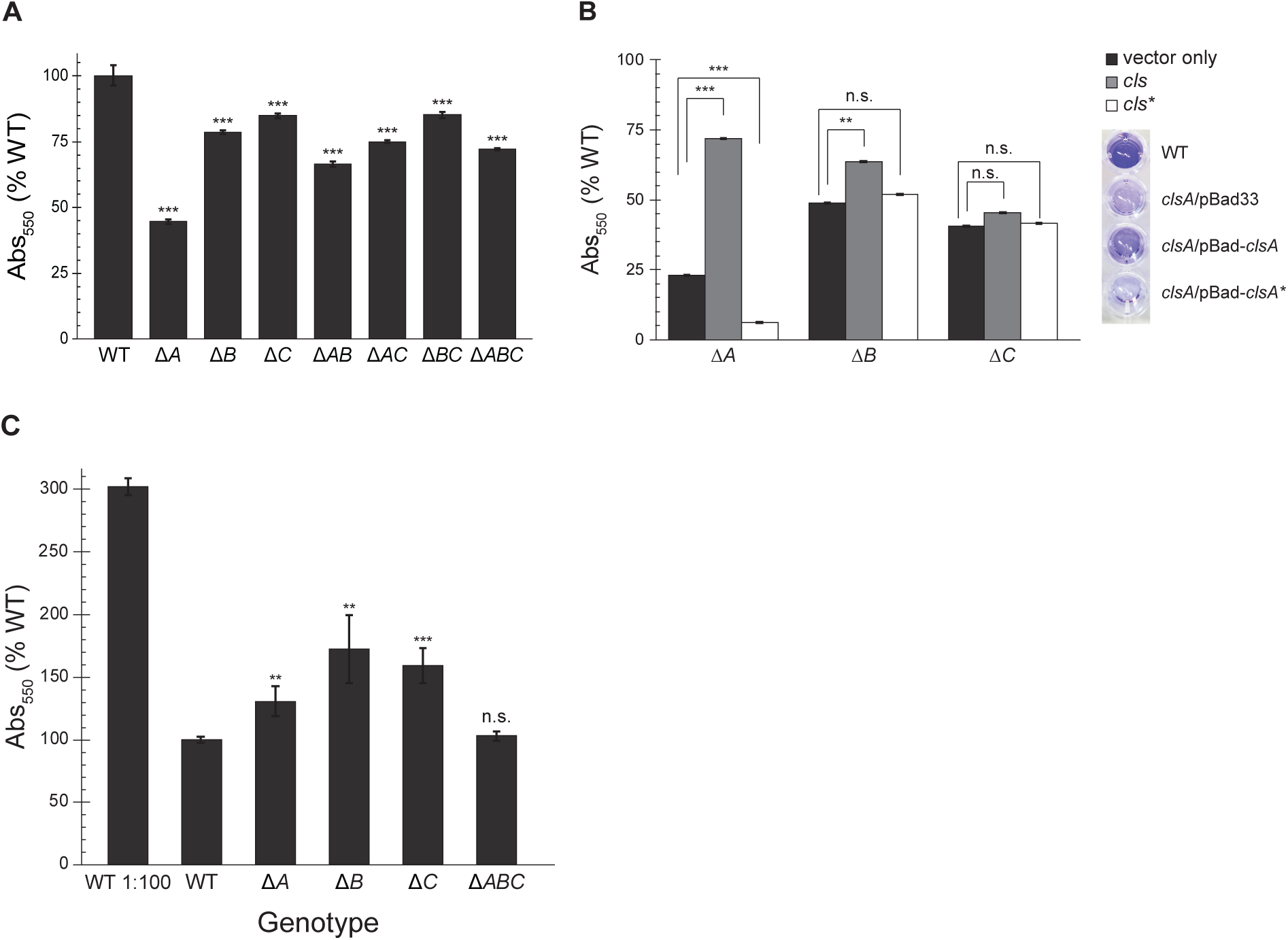
Disruption of cardiolipin synthesis reduces biofilm formation. *E. coli* cells were grown in microtiter plates for 24 h at 30°C without shaking. Adherent cells were stained with crystal violet (CV), and CV absorbance was measured (λ=550 nm). Error bars indicate standard error; \*\*\**p* < 0.001, \*\**p* < 0.01, \**p* < 0.05, n.s. not significant; Student’s *t* test compared to WT. (A) CV labeling of mutant *E. coli* cells with noted genotypes. (B) Δ*cls* strains were complemented with pBad33 (vector), full-length *cls*, or a catalytically inactive mutant (*cls\**; *clsA* H224F H404F, *clsB* H113A, or *clsC* H130A). Cells were grown with 0.2% arabinose to induce protein expression; CV absorbance values of cells grown with arabinose were normalized to CV absorbance of uninduced cells. Error bars indicate standard error; \*\*\**p* < 0.001, \*\**p* < 0.01, n.s. not significant; Student’s *t* test compared to uninduced control. Inset: image of representative crystal violet staining of cells grown in M9 with 0.2% arabinose. From top: wild-type (MG1655); Δ*clsA*/pBad33; Δ*clsA*/pBad-clsA; Δ*clsA*/pBad-clsA*. (C) CV staining of biofilms started using saturated overnight cultures; CV labeling of biofilm started with subcultured WT is shown for reference. Error bars indicate standard error; \*\*\**p* < 0.001, \*\**p* < 0.01, n.s. not significant; Student’s *t* test compared to WT.

While ClsA is active at all stages of growth, ClsB and ClsC do not contribute significantly to CL production until stationary phase or under certain conditions of stress (23, 34). We noted that Δ*clsA* mutants exhibited a more drastic reduction in surface attachment than other *cls* deletion strains. To investigate the growth phase dependence of surface attachment in *cls* mutants, we assayed the ability of stationary phase cells to adhere to surfaces. We used saturated overnight cultures to inoculate 96-well microplates and measured crystal violet labeling after 24 h of incubation (Fig. 3*C*). Under these conditions, strains lacking a single *cls* gene had a higher number of cells adhered to microplate well surfaces than a mutant completely lacking CL.

Deletion of *clsA* has been associated with increased sensitivity of cells to novobiocin, an aminocoumarin-based inhibitor of bacterial DNA gyrase (35). As the antibiotic sensitivity may arise by altering the structure of the IM and increasing the transport of novobiocin across the membrane, we explored whether cells lacking CL were more susceptible to other families of antibiotics—including those targeting the membrane. We determined the minimum inhibitory concentrations (MIC) of multiple classes of antimicrobial compounds, including two targeting the cell membrane: cecropin A (an antimicrobial peptide), and polymyxin B (a mixture of lipopeptides derived from the bacterium *Bacillus polymyxa*) (Table *S*4). In general, planktonic *E. coli* Δ*clsABC* cells were more susceptible to these compounds relative to surface-attached cells. Surprisingly, treatment of both WT and CL-deficient cells attached to surfaces with polymyxin B or cecropin A slightly *increased* attachment compared to a non-treated control (Fig. *S*4), but none of the antimicrobials we tested showed a significant difference in their effects on surface attachment between cells of WT and *cls* mutant strains.

### Cardiolipin impacts activation of the Rcs envelope stress response

Depletion of PE or PG causes severe physiological defects, decreases cell viability, and activates multiple stress responses; in contrast, depletion of CL has little quantifiable effect on cell physiology (12). Previous studies largely focused on the effects of disrupting ClsA. We reasoned that removing CL completely by deleting all three CL synthases would likely have a more dramatic physiological effect and that activation of membrane stress response(s) was a potential link between CL depletion and biofilm defects.

*E. coli* possesses five signaling pathways known to be associated with envelope stress: Psp (<u>p</u>hage <u>s</u>hock <u>p</u>rotein), Bae (<u>b</u>acterial <u>a</u>daptive r<u>e</u>sponse), σ^E^, Rcs (<u>r</u>egulation of <u>c</u>olanic acid <u>s</u>ynthesis), and Cpx (<u>c</u>onjugative <u>p</u>ilus e<u>x</u>pression) (36). We measured the activation of these pathways using quantitative PCR (qPCR) of genes transcriptionally regulated by each specific signal transduction pathway (Fig. 4*A*). Rcs-activated transcripts were on average 20-fold more abundant in Δ*clsABC* cells than in WT cells. Further, disruption of the Rcs pathway was sufficient to restore the attachment of cells to WT levels in a surface attachment assay (Fig. 4*B*).

**Figure 4.**
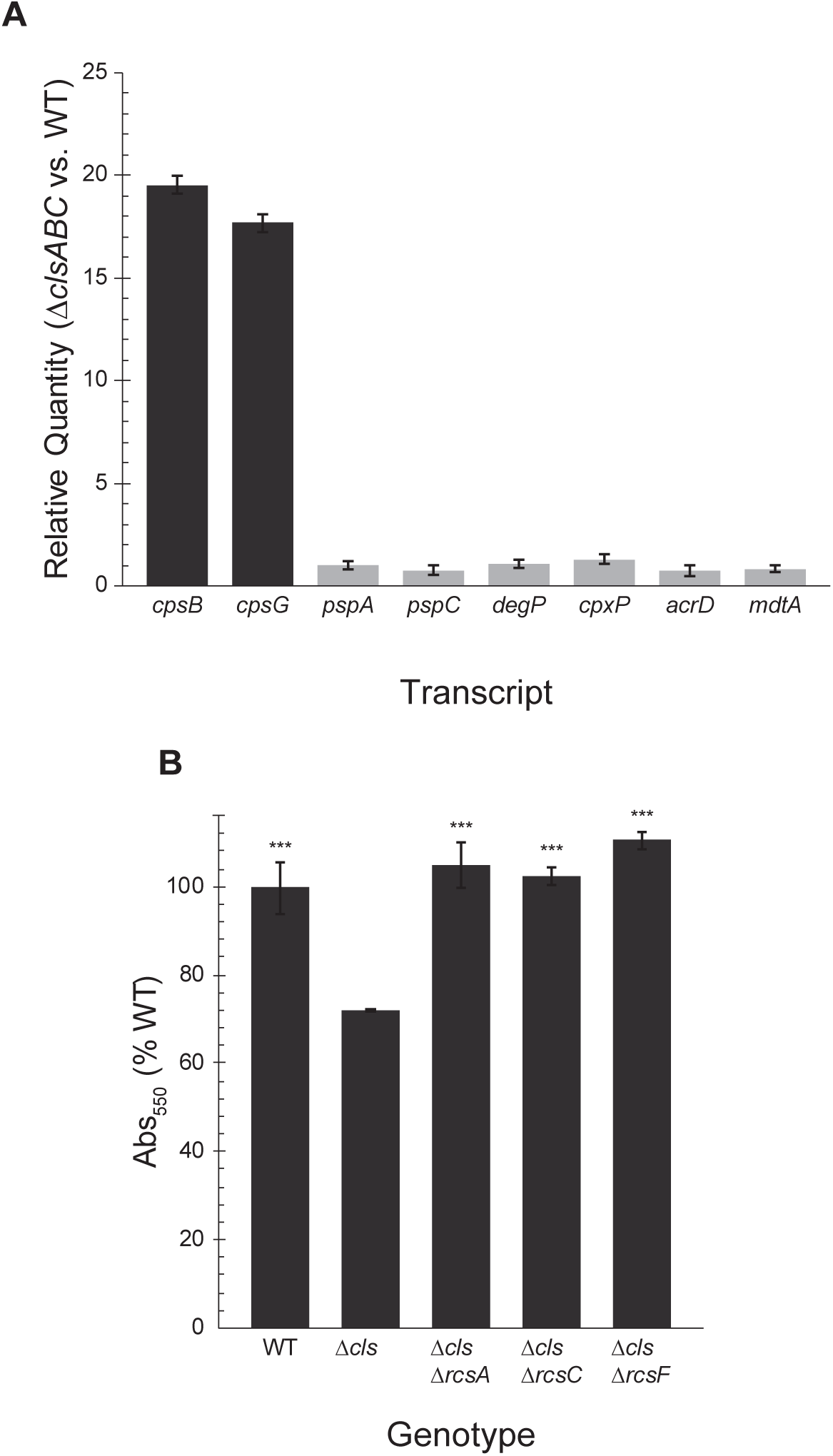
The Rcs phosphorelay is activated in CL deficient cells. (A) We used qPCR to measure activation of 5 major envelope stress pathways in Δ*clsABC* (Δ*cls*) cells. The relative abundance of transcripts of the Rcs-controlled genes *cpsB* and *cpsG* was ∼20-fold higher in Δ*cls* compared to wild-type MG1655 cells. Bae (*mdtA, acrD*), Psp (*pspA, pspC*), sigma E (degP), and Cpx (*cpxP, degP*) signaling pathways were not significantly activated (0.5 ≤ RQ ≤ 2). Error bars indicate standard deviation. (B) Crystal violet staining of Δ*cls* mutants, and Δ*rcs* mutants in a Δ*cls* background. Error bars indicate standard error; \*\*\**p* < 0.001; Student’s *t* test compared to Δ*cls*.

Initiation of the Rcs signaling system causes the phosphorylation of RcsA and RcsB, enabling them to function as transcriptional regulators (Fig. 5). RcsB (which can form a homodimer in its phosphorylated state) and RcsAB regulate a diverse array of genes. The Rcs regulon includes a number of genes involved in acid resistance, as well as the colanic acid and curli synthesis operons, osmotically inducible peroxidase *osmC*, and the flagellar master regulator *flhDC* (28). Production of colanic acid and other secreted polysaccharides promotes later stages of biofilm maturation by providing structure and protection to the developing community. Improperly timed activation of Rcs impairs the functioning of other systems required for biofilm formation, including expression of flagella (37-39). Further, PG deficient *E. coli* cells exhibit defects in flagella synthesis (12, 40). By immunostaining cells with an antibody raised against FliC, we observed cells lacking CL had significantly fewer flagella than WT cells (Fig. 6*A* and *B*) and displayed a decrease in both swimming and swarming motility (Fig. 6*C* and *D*). Deleting the RcsA transcriptional regulator, which binds phosphorylated RcsB to form a complex that represses *flhDC*, increased production of flagella in the *cls* mutants and rescued swarming motility defects.

**Figure 5.**
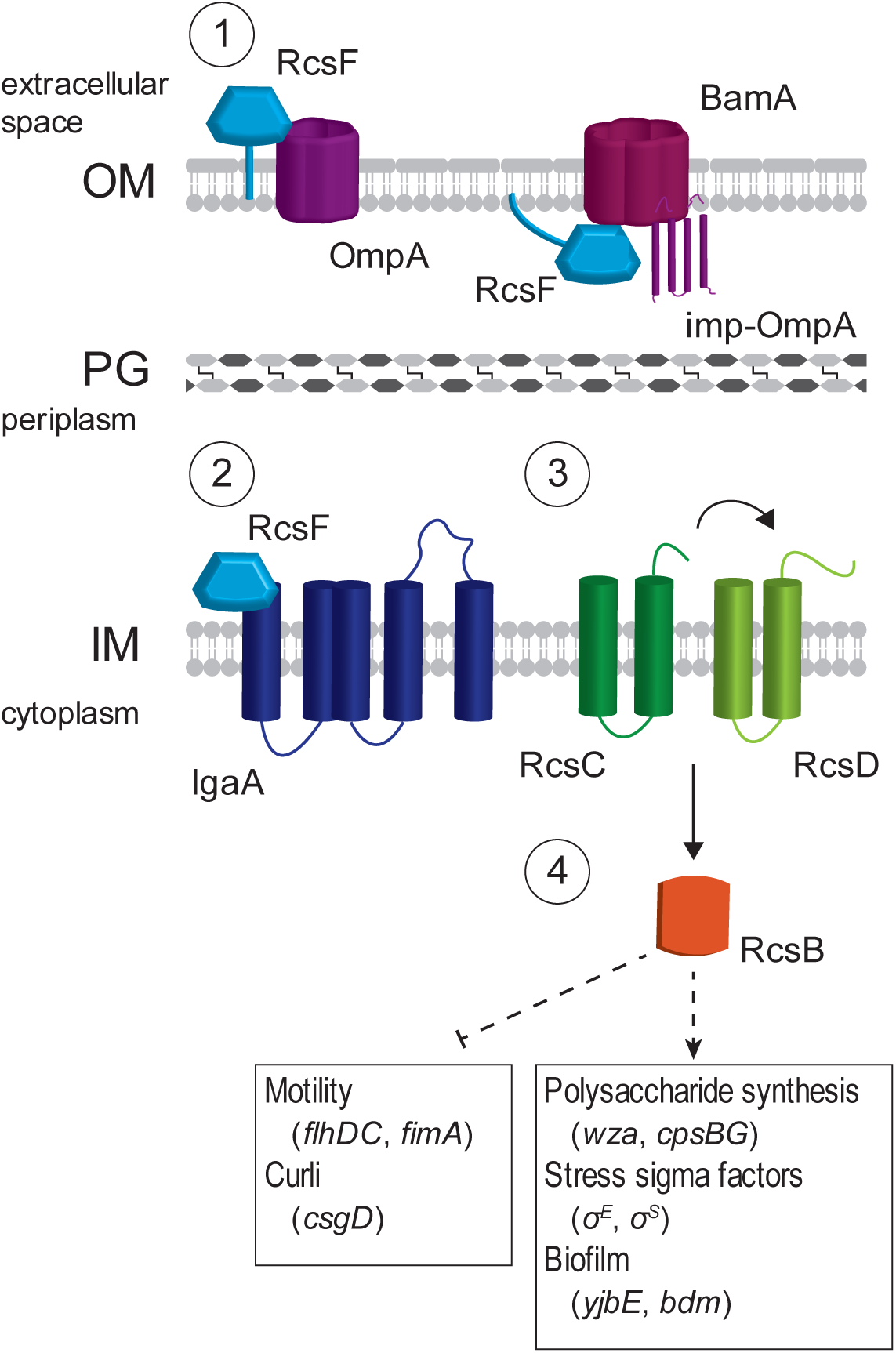
Rcs monitors export of outer membrane proteins. (1) After its synthesis, RcsF is transported to the outer membrane by the chaperone LolA and associates with BamA, a component of the machinery that assembles β-barrel proteins; the immature, processed form of OmpA is referred to as ‘imp-OmpA’. RcsF is then assembled into a complex with OmpA. (2) Under normal conditions, IgaA represses the Rcs phosphorelay. However, defects in protein translocation cause a buildup of free RcsF, which associates with IgaA and blocks its inhibition of the phosphorelay. (3) Once repression by IgaA is released, RcsC is autophosphorylated. The phosphate is transferred to RcsD, then to the transcription factor RcsB. (4) Phosphorylated RcsB interacts with itself and other transcription factors to regulate genes involved in a number of biofilm formation processes, as well as genes for acid resistance, cell division, and other transcriptional regulators.

**Figure 6.**
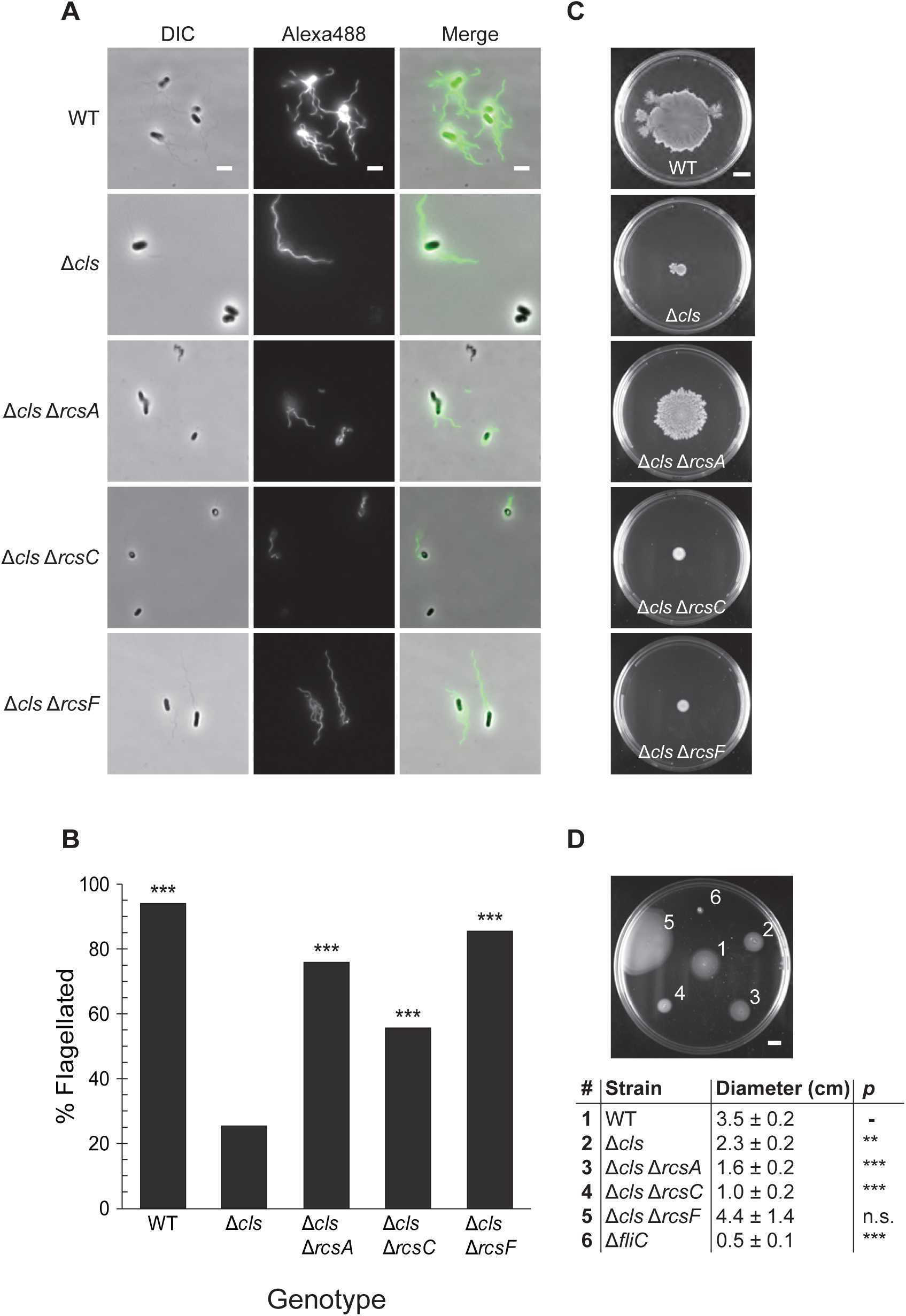
Cardiolipin deficient cells produce fewer organelles that enable surface attachment and promote biofilm formation. (A) Using an antibody raised against FliC, we immunostained cells in the late exponential phase of growth and (B) determined the percentage of cells with at least one visible flagellum (N ≥ 100 cells). DIC = differential interference contrast. \*\*\**p* < 0.001; Fisher’s exact test, compared to Δ*clsABC*. Scale bar is 3μm. (C) Swarm and (D) swim plates were inoculated as described in the Materials and Methods, and imaged after 24 h at 30°C. Average swimming diameters after 24 h and standard deviations are indicated for each strain. \*\**p* < 0.01, \*\*\**p* < 0.001, n.s. not significant; Student’s *t* test compared to WT. Scale bar is 1 cm.

### Cardiolipin enhances protein translocation *in vivo*

RcsF is an outer membrane lipoprotein that relays stress signals to the sensor kinase RcsC (Fig. 5). Newly synthesized RcsF is transported to the outer membrane (OM) β-barrel assembly complex by the periplasmic chaperone LolA, where it is assembled into a complex with OmpA (41, 42). This step sequesters RcsF and prevents it from binding IgaA, an IM protein that downregulates the Rcs pathway (43). Defects in the maturation of OM proteins increases the size of the pool of unbound RcsF that is able to bind IgaA, thus relieving the inhibition of Rcs activation by IgaA.

The majority of periplasmic and membrane-associated proteins are translocated across the IM by the Sec translocon—a large, multi-subunit complex that is stimulated by CL when reconstituted in proteoliposomes (44, 45). We hypothesized that depleting CL would have a negative effect on Sec-mediated protein translocation *in vivo*. As a reporter system to assay protein translocation activity in *clsABC* mutants, we tracked translocation of the alkaline phosphatase PhoA, which is inactive until it is exported to the periplasm (Fig. 7). WT cells exported PhoA at a rate of 20.3 ± 0.7 units/h. In contrast, we found that cells lacking CL had a significantly decreased level of PhoA translocation activity (2.9 ± 0.5 units/h) that was comparable to the translocation rate of a Δ*secG* strain (3.1 ± 0.4 U/hr) (Fig. 7). Further, Western blot analysis of His-tagged OmpA showed that WT cells produced mature OmpA at a faster rate than Δ*clsABC* cells (Fig. *S*5).

**Figure 7.**
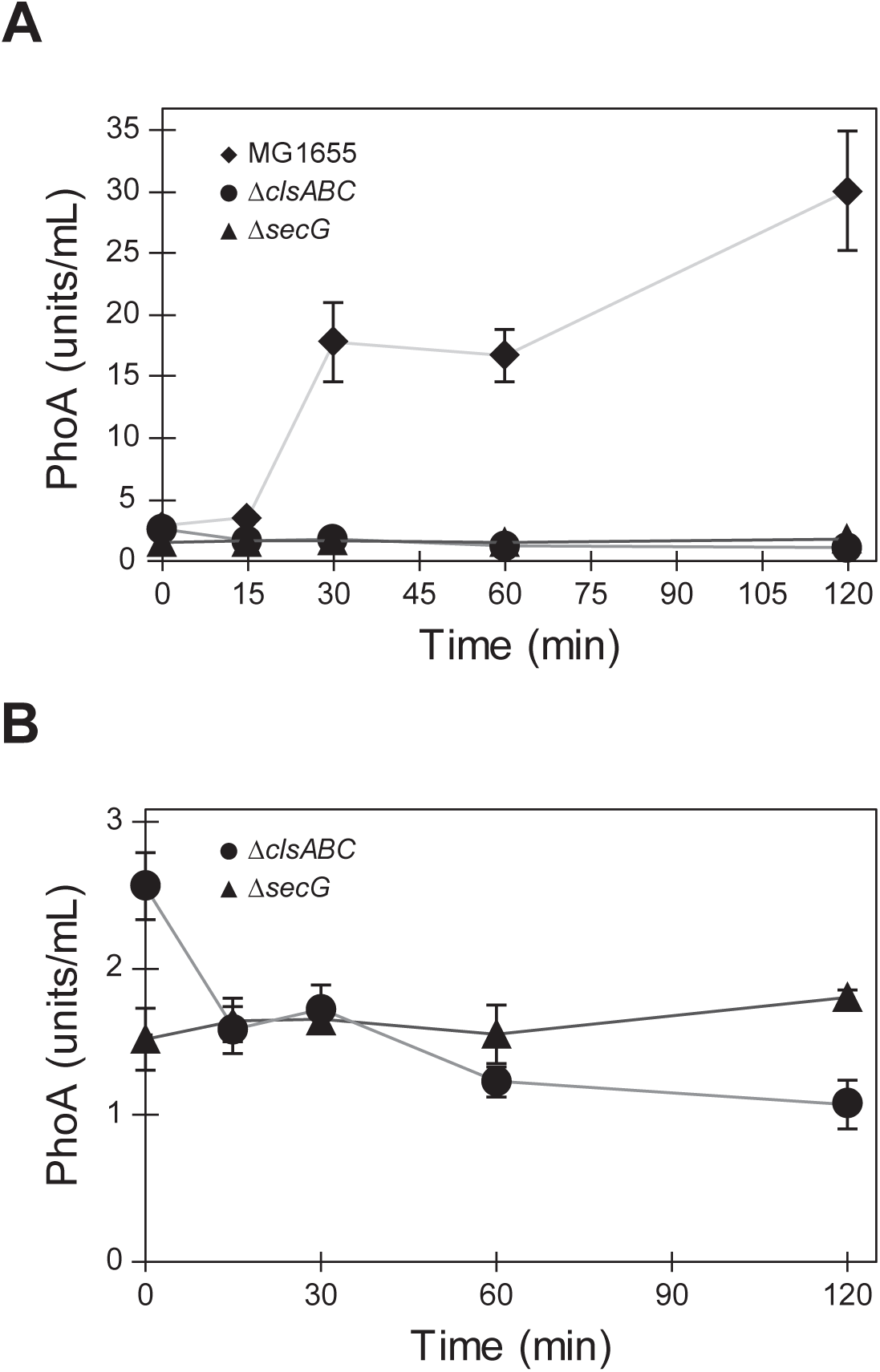
Protein translocation is reduced in the absence of CL. (A) and (B) Periplasmic activity of PhoA in wild-type MG1655 (diamonds), Δ*clsABC* (circles), and Δ*secG* (triangles); 1 unit = 1 mol/min of hydrolyzed PNPP. Measurements represent the average of 3 biological replicates and error bars indicate standard error of the mean.

## Discussion

The physiological roles of CL are still emerging and *E. coli* has played a key role as a model bacterium for understanding this family of phospholipids. Various CL-protein interactions have been identified, including interactions with respiratory complexes, aquaporins, and DNA recombination proteins (e.g., RecA and DnaA) (26, 46-49). The CL composition in cells is correlated with entry into stationary phase and osmotic stress (23, 50, 51). A growing body of evidence suggests that CL stabilizes cell membranes at regions of high cell wall curvature—e.g., at sites of cell division and spore formation—by relieving elastic strain through the reorganization of phospholipids within the liquid crystalline bilayer (52, 53).

CL synthases are redundant and numerous studies have demonstrated that they are non-essential. It is likely that PG supports some of the cellular function of CL when the latter is missing in the membrane; significant cell defects arise when both CL and PG are removed from cells, and cell viability in these mutants requires accessory mutations in Braun’s lipoprotein, Lpp. In our biofilm studies, it appears that PG is not sufficient to replace the change in membrane properties that arise from depleting CL.

Many previous studies of CL centered on *clsA* mutants and almost all studies of CL and CL synthases have been performed using cells growing in optimized laboratory conditions. We studied the effects of CL depletion on processes involved in biofilm development, a lifestyle commonly adopted by cells under stress that can be considered very from the ideal laboratory growth conditions referred to above, including starvation and deprivation of specific growth factors. Many studies of CL synthases in *E. coli* have been conducted using the rich medium, lysogeny broth (LB), however enteric bacteria are likely to encounter fluctuating environments that are not well mimicked by persistently high concentrations of nutrients; instead these cells likely experience short bursts of high concentrations of nutrients interspersed with long periods of low nutrient concentrations (54). In minimal nutrient medium, we identified a biofilm phenotype associated with a cellular CL deficiency, which was not observed with cells grown in LB media (Fig. *S*2). Consequently, a biofilm phenotype arising from the absence of CL in cells growing in low nutrient conditions is consistent with previous observations that biofilms are sensitive to the environmental context in which they are grown (55-57).

We found that Δ*clsA* cells showed a more severe surface attachment defect than other *cls* mutants, including combinations of Δ*clsA* and Δ*clsB* and/or Δ*clsC* (Fig. 3*A*). This phenotype may be partially due to differential activity of the synthases at different stages of cell growth; i.e., ClsA appears to be constitutively active, while ClsB and ClsC have low activity until stationary phase (23). To test this hypothesis, we assayed the earliest stage of biofilm formation—i.e., attachment of cells to a surface—starting from saturated cultures; a point in time at which all 3 synthases contribute to CL production (Fig. 3*C*). In this context, single *cls* deletion mutants attach more effectively than a strain that lacks all three cardiolipin synthases, supporting our initial assumption. However, the observation that strains lacking multiple synthases attach better and produce more biofilm than cells missing only *clsA* suggests that the interplay between these enzymes is more complex than a simple additive effect.

We demonstrated that the Rcs phosphorelay system is activated in CL-deficient mutants growing in nutrient-limited conditions and impairs biofilm formation. We found that CL-deficient mutants have fewer flagella and reduced swimming and swarming cell motility compared to WT cells. These observations are in agreement with Rcs activation causing decreased expression of *flhDC* and reduced motility in other gram-negative enteric bacteria, such as *Serratia marcescens* and *Salmonella enterica* (58, 59). Disrupting the transcription of components of the Rcs signal transduction pathway in *E. coli* restored surface attachment to WT levels and improved flagella production. Deletion of *rcsF* resulted in the greatest improvement in flagella production and swimming migration, however Δ*rcsA* was the only mutant to display a significant improvement in swarming, suggesting that Rcs activity affects relevant factors other than the number of flagella in *cls* mutants. *E. coli* mutants lacking *rcsC* performed better than Δ*clsABC* only in surface attachment assays. Consistent with these findings, Rcs activity has been reported as an important step in late stages of biofilm formation, yet alterations in the timing of Rcs pathway activation inhibit organelle production and reduce the ability of cells to attach to surfaces (37, 39).

Rcs activity also influences the formation of fimbriae in *E. coli* by affecting the transcription of *fimB* and *fimE* (60). The relative ratio of the FimB and FimE recombinases dictates the orientation of the *fimA* promoter (*fimAp*), and thus controls transcription of the entire type I fimbriae operon (61). We did not observe any change in the ratio of phase-on (*fimAp* facing the *fim* operon) vs. phase-off (*fimAp* facing opposite to *fim* operon, preventing transcription) oriented cells in the absence of *clsABC* (Fig. *S*6); however, it is possible that an investigation of the number of fimbriae per cell may tell a different story.

Rowlett *et al.* (2017) described the activation of multiple envelope stress pathways in the absence of CL using fluorescent protein promoter fusions and immunoblotting to measure protein levels; we found no significant activation of pathways other than Rcs using qPCR to directly quantify transcript levels of target genes (22). As each of these techniques interrogates different steps in protein expression, our results do not contradict those published previously. A detailed investigation of pre- and post-transcriptional changes in CL-deficient cells would help further illuminate potential factors affecting biofilm development in CL mutants.

The activity of stress pathways such as Rcs has widespread effects on cell physiology: e.g., *rcsB* overexpression affected transcript levels of multiple genes involved in drug efflux and resistance (62). To test whether cls mutants may yield synthetic lethal phenotypes when matched with the loss of function of other proteins, including those involved in the structure and function of components of the cell wall, we paired the effects of CL depletion with different antibiotics. We observed that planktonic *E. coli* Δ*clsABC* cells had an increased susceptibility to several antimicrobial agents (Table *S*4), which supports this hypothesis. We also tested whether these antibiotics reduced biofilm formation (including cell attachment) of planktonic *E. coli* Δ*clsABC* cells and found the effects variable and generally inconclusive (Fig. *S*3).

Anionic phospholipids and Rcs activation have been discussed together in multiple studies with limited biochemical data points to support their connection. Previous studies of CL and the Sec translocon suggested a physical link between these two cellular components that may lead to Rcs activation (44). CL is tightly bound to purified, recombinant SecYEG and has been reported to enhance the stability of the SecYEG complex and stimulate the ATPase activity of SecA *in vitro*. SecYEG substrates include lipoproteins, whose translocation and assembly are monitored by the outer membrane lipoprotein RcsF (41). The observation of regions of SecYEG to which CL binds suggests why PG may not be able to replace the loss of CL impacting the function of protein translocation. Accordingly, our results demonstrate that mutants deficient in CL display a dramatic reduction in protein translocation activity, supporting a role for SecYEG in the interaction between CL and Rcs. Other work has shown that the absence of CL causes an increase in unfolded OmpF, further supporting our hypothesis (22).

The Rcs phosphorelay system consists of integral membrane proteins that may have activities that respond to changes in membrane fluidity, which occurs with other classes of membrane proteins (63-65). Increasing the concentration of CL increases membrane fluidity *in vitro*, however the impact of depleting CL on the fluidity of membranes in living cells has not yet been demonstrated conclusively (66). A recent study found no significant reduction in membrane fluidity in Δ*clsA E. coli* cells using fluorescence recovery after photobleaching (FRAP) experiments of membranes doped with BODIPY-labeled C-12 fatty acids (67). This work did not examine fluidity in a strain that was entirely CL-null, nor did it investigate the effect of multiple growth conditions. FRAP is generally unable to detect small changes in cell membrane fluidity, and the structure of the bacterial membrane in live cells—containing large amounts of lipopolysaccharide—may confound the interpretation of data.

Several studies have demonstrated that saturated fatty acids (SFAs) inhibit swarming in multiple organisms, such as *S. enterica*, a pathogen closely related to *E. coli*. As swarming is reported to be a stage connected to biofilm formation, we hypothesize that the presence of SFAs—and thus membrane fluidity—may also play a role in biofilm formation (19, 68). A recent study demonstrated that cells of *Staphylococcus aureus, L. monocytogenes, P. aeruginosa*, and *S. Typhimurium* in biofilms are enriched in SFAs compared to planktonic cells (69). Our studies focus on the effects of phospholipid composition of cells in relatively early stages of biofilm development. A more detailed investigation of biofilm membrane composition that includes an examination of various types of biofilms (e.g. flow cell and pellicle) at different stages of development, and analyzes changes in lipids and the composition of their acyl tail groups may reveal details on the interplay of lipids and biofilm formation in *E. coli*.

Several outstanding questions concerning the role of CL in cellular processes remain unanswered. First, disruption of CL synthesis has broad effects on transcriptional regulation of a variety of genes, however the biochemical mechanisms that regulate production of CL are unknown. Second, a better understanding of CL regulation may aid in discovering the specific function(s) of each individual CL synthase and understanding the purpose of their redundancy in *E. coli*. The presence of multiple CLS genes is widely conserved among bacteria, with most species encoding at least 2 synthases. A BLAST search reveals that the closely related bacterium *S. enterica* encodes for proteins with 94%, 87%, and 82% identity to *E. coli* ClsA, ClsB, and ClsC, respectively (70). Available data points to a hypothesis in which CL serves a variety of physiological functions. The results described in this paper expands our understanding of the roles of CL and other membrane lipids in biofilms in conditions that mimic the natural environment of *E. coli* cells.

## Materials and Methods

### Strains and growth conditions

All bacterial strains used in this study are listed in Table S1 (Supplemental Information). We created mutants using protocols for P1 phage transduction, chemical transformations, and lambda-red recombination (71-73). To construct strains of *E. coli*, we grew cells at 37°C in lysogeny broth (LB) consisting of 1% [w/v] tryptone, 0.5% [w/v] yeast extract, and 1% [w/v] NaCl, or on LB plates containing 1.5% agar and infused with LB. For all other purposes, cells grew at 30°C in M9 minimal media (3.4% [w/v] Na2HPO_4_, 1.5% [w/v] KH_2_PO_4_, 0.25% [w/v] NaCl, 0.5% [w/v] NH_4_Cl, 0.05% [w/v] thiamine HCl, 2 mM MgSO_4_, 0.1 mM CaCl_2_, 0.4% [w/v] glucose) supplemented with a defined amino acid mixture (500 mg/mL of each alanine, cysteine, glycine, histidine, aspartic acid, glutamic acid, phenylalanine, asparagine, glutamine, methionine, leucine, isoleucine, proline, serine, threonine, lysine, and valine, 50 mg/mL tryptophan, and 50 mg/mL tyrosine). Antibiotics (50 μg/mL ampicillin, 20 μg/mL tetracycline, 30 μg/mL kanamycin, and/or 25 μg/mL chloramphenicol) were added to growth media as needed. To induce expression of genes from various plasmid constructs, we added L-arabinose to a final concentration of 0.2% (w/v).

### Surface attachment assays

We quantified early stages of *E. coli* biofilm formation using a crystal violet assay by diluting cells from an overnight M9 culture 1:100 into fresh M9 in wells of polystyrene 96-well microplates (100 μL/well), and incubating at 30°C for 24 h unless indicated otherwise (74). We measured optical density by quantifying absorbance of the wells at λ=600 nm (OD_600_). Liquid growth media was removed and discarded, plates were rinsed with distilled water to remove non-adherent cells, 125 μL of an aqueous solution of 0.1% crystal violet was added to each well, and plates were incubated for 15 min at 25°C. The liquid in each well was removed and wells were rinsed with distilled water 3 times; addition of 125 μL of 30% acetic acid (in water) released crystal violet trapped in biofilms, and the dye was quantified by measuring the absorbance at λ=550 nm. We normalized the absorbance of crystal violet to absorbance of cells in the starting culture (OD_600_) and determined the statistical significance of the data using an unpaired *t* test.

### Quantitative polymerase chain reaction

We used a Zymo Research Direct-zol RNA Miniprep kit (Zymo Research Corp., CA, USA) to extract total RNA from *E. coli* cells. Genomic DNA was removed using the ArcticZymes HL-dsDNase (ArcticZymes, Norway), and RNA was reverse transcribed using an Applied Biosystems High Capacity RNA-to-cDNA kit (Life Technologies, TX, USA). We treated newly synthesized cDNA with RNase H (New England Biolabs, MA, USA) to digest RNA hybridized to cDNA. A PowerUp SYBR Green Master Mix (Life Technologies, TX, USA) enabled us to perform quantitative PCR on an Applied Biosystems 7500 Fast Real-Time PCR System (Applied Biosystems, CA, USA) using the manufacturer’s instructions for a standard cycling protocol. We used primers for *gapA* and/or *idnT* as endogenous controls.

### Motility assays

For *E. coli* swarming assays, we prepared swarm plates by pipetting 15 mL of hot LB containing 0.6% Eiken agar (Japan) and 0.5% [w/v] glucose into 100 x 15-mm Petri dishes (BD). We cooled plates uncovered for 30 min in a laminar flow hood, then inoculated the center of each plate with 3 μL of a saturated *E. coli* culture (∼10^9^ cells/mL). We waited <5 min for the excess liquid in the droplet to absorb into the agar, covered the plates, and incubated the inverted plates at 30°C for 24 h. For swimming assays, we prepared plates using M9 minimal media, without amino acids, containing 0.3% agar. We allowed the plates to solidify and cool uncovered for 30 min in a laminar flow hood, then inoculated with 3 μL of a saturated *E. coli* culture. After drying for <5 min, the plates were incubated, right-side up, at 30°C for 24 h.

### Immunofluorescence microscopy of flagella

Immunostaining of flagella was performed using an anti-FliC primary antibody and an AlexaFluor 488-tagged goat anti-rabbit IgG secondary antibody using a published protocol (68). We grew *E. coli* cells to an absorbance (λ=600 nm) of 0.6-0.8, diluted cells 1:5 in 1X PBS, and imaged them in chambers consisting of a glass coverslip attached to a glass slide with double-sided tape and treated with a solution of poly-L-lysine. After adding cells to the chambers, we fixed them with 1% formaldehyde in PBS then rinsed and incubated cells in blocking buffer [3% bovine serum albumin (BSA) containing 0.2% Triton X-100 in 1X PBS] overnight at 4°C. We incubated *E. coli* cells in a solution of primary antibody diluted 1:1,000 in blocking buffer for 1 h at 25°C, washed with washing buffer (0.2% BSA and 0.05% Triton X-100 in 1X PBS), then incubated in a solution of secondary antibody for 1h. We incubated cells in secondary antibody diluted 1:1,000 in blocking buffer for 1 h, washed with washing buffer followed by PBS, and imaged using a Nikon TI-E Eclipse inverted epifluorescence microscope.

### PhoA translocation assay

We measured PhoA translocation as described previously (75). Overnight cultures of *E. coli* were diluted 1:50 in fresh M9 and grown to an absorbance of 0.6-0.8 (λ=600). We induced the expression of PhoA in *E. coli* cells from pBad33 using 0.2% arabinose, removed 1 mL of culture at the indicated timepoints for analysis, and determined the absorbance (λ=600). We arrested protein secretion by addition of iodoacetamide to a final concentration of 2 mM, collected cells by centrifugation for 1 min at 12,000 × *g* at 4°C, discarded the supernatant, and resuspended cells in 1 mL MOPS buffer (67 mM 3-(*N*-morpholino)propanesulfonic acid, 83 mM NaCl, 16 mM NH_4_Cl, 10 mM MgCl_2_, pH 7.2). We repeated the wash step once more, then added 100 μL of the *E. coli* cell suspension to a microcentrifuge tube containing 900 μL TZ buffer (1 M Tris-HCl, 1 mM ZnCl_2_, pH 8.1), 25 μL 0.1% [w/v] SDS, and 25 μL of CHCl_3_. Cells suspensions were vortexed briefly, then 100 μL of 4 mg/mL *p*-nitrophenyl phosphate (New England Biolabs, MA, USA) was added and the mixture was vortexed again. Assay tubes were incubated at 28°C until the solution began to turn yellow, then centrifuged for 5 min at 12,000 × *g* at 4°C. We removed 800 μL of liquid from the upper portion of the tube, measured the absorbance of this aliquot (λ=420 nm), and calculated units of active PhoA using the equation: (1000 × A_420_)/(*t* × V × OD_600_), where *t* is the time after induction of PhoA expression in min and V is the volume of cell suspension added in mL.

### Lipid extraction and thin layer chromatography

We extracted and quantified total membrane lipids from *E. coli* cells using a modified version of the Bligh and Dyer method (76). Briefly, we pelleted *E. coli* cells at 5,000 x *g* for 10 min, resuspended in 100 μL of water, and lysed cells by adding 125 μL CHCl_3_ and 250 μL methanol and inverting tubes several times. We added 100 μL of H2O and 100 μL CHCl_3_, vortexed samples, centrifuged for 5 min at 13,000 rpm, collected the lower organic phase, and dried it under a stream of N2. We dissolved dried lipid extracts in 40 μL of 1:1 methanol:CHCl_3_ and used 5-10 μL of sample for thin layer chromatography (TLC) on TLC silica gel 60 plates (Merck, Germany) with a mixture of 65:25:10 CHCl_3_:CH_3_OH:CH_3_COOH for the mobile phase. After drying, we sprayed TLC plates with a cupric sulfate solution (100 mg/mL CuSO_4_ in 8% [w/v] H_3_PO_4_) and incubated TLC plates on heat plates at 145°C for 5 min. We imaged plates using the Cy3 fluorescence setting on a GE ImageQuant LAS 4010 (GE Healthcare Bio-Sciences, PA, USA) and quantified the intensity of bands using Image J version 1.51h.

To extract lipids from biofilm-associated *E. coli* cells, we grew cells in 96-well microplates as described. After discarding the waste media and rinsing the wells of the plates, we added 50 μL of 1.5 M NaCl to each well and detached *E. coli* cells from the microplate surface by sonicating for 10 min in a Branson 2510 bath sonicator (Branson, CT, USA). We collected resuspended *E. coli* cells, centrifuged them for 10 min at 5,000 × *g*, removed the supernatant, and proceeded with lipid extraction and quantification as described above.

### Statistical analysis

We used a Student’s *t* test for pair-wise comparisons and a Fisher’s exact test for comparisons of categorical variables. We performed experiments with at least 3 independent biological replicates to ensure reproducibility. We considered *p* values less than 0.05 statistically significant. For qPCR experiments, we considered changes of at least twofold to be significant.

## Acknowledgements

We acknowledge funding from NSF (pre-doctoral fellowship DGE-1256259 to J.F.N.; grant DMR-1121288) and the USDA (grants WIS01594 and WIS01942). We are grateful to Howard Berg’s lab for providing the anti-FliC antibody.

## References

1. Davey ME, O’Toole GA. 2000. Microbial biofilms: from ecology to molecular genetics. Microbiol Mol Biol Rev 64:21.

2. Hall-Stoodley L, Costerton JW, Stoodley P. 2004. Bacterial biofilms: from the natural environment to infectious diseases. Nat Rev Microbiol 2:95–108.

3. Woo KY, Keast D, Parsons N, Sibbald RG, Mittmann N. 2015. The cost of wound debridement: a Canadian perspective. Int Wound J 12:402–407.

4. Schultz MP, Bendick JA, Holm ER, Hertel WM. 2011. Economic impact of biofouling on a naval surface ship. Biofouling 27:87–98.

5. Campoccia D, Montanaro L, Arciola CR. 2013. A review of the clinical implications of anti-infective biomaterials and infection-resistant surfaces. Biomaterials 34:8018–8029.

6. Kostakioti M, Hadjifrangiskou M, Hultgren SJ. 2013. Bacterial biofilms: development, dispersal, and therapeutic strategies in the dawn of the postantibiotic era. Cold Spring Harb Perspect Med 3:a010306.

7. Pratt LA, Kolter R. 1998. Genetic analysis of *Escherichia coli* biofilm formation: roles of flagella, motility, chemotaxis and type I pili. Mol Microbiol 30:9.

8. O’Toole G, Kaplan HB, Kolter R. 2000. Biofilm formation as microbial development. Annu Rev Microbiol 54:49–79.

9. Serra DO, Hengge R. 2014. Stress responses go three-dimensional—the spatial order of physiological differentiation in bacterial macrocolony biofilms. Environ Microbiol 16:17.

10. Flemming H-C, Wingender J. 2010. The biofilm matrix. Nat Rev Microbiol 8:11.

11. Danese PN, Pratt LA, Kolter R. 2000. Exopolysaccharide production is required for development of *Escherichia coli* K-12 biofilm architecture. J Bacteriol 182:4.

12. Uchiyama J, Nobue Y, Zhao H, Matsuzaki H, Nagahama H, Matsuoka S, Matsumoto K, Hara H. 2010. Involvement of sigmaS accumulation in repression of the flhDC operon in acidic phospholipid-deficient mutants of Escherichia coli. Microbiology 156:1650–1660.

13. Uchiyama J, Sasaki Y, Hagahama H, Itou A, Matsuoka S, Matsumoto K, Hara H. 2010. Accumulation of sS due to enhanced synthesis and decreased degradation in acidic phospholipid-deficient *Escherichia coli* cells. FEMS Microbiol Lett 307:8.

14. Beloin C, Valle J, Latour-Lambert P, Faure P, Kzreminski M, Balestrino D, Haagensen JAJ, Molin S, Prensier G, Arbeille B, Ghigo J-M. 2003. Global impact of mature biofilm lifestyle on Escherichia coli K-12 gene expression. Molecular Microbiology 51:659–674.

15. Dorel C, Vidal O, Prigent-Combaret C, Vallet I, Lejeune P. 1999. Involvement of the Cpx signal transduction pathway of E. coli in biofilm formation. FEMS Microbiol Lett 178:169–175.

16. Benamara H, Rihouey C, Jouenne T, Alexandre S. 2011. Impact of the biofilm mode of growth on the inner membrane phospholipid composition and lipid domains in Pseudomonas aeruginosa. Biochim Biophys Acta 1808:98–105.

17. Benamara H, Rihouey C, Abbes I, Mlouka MAB, Hardouin J, Jouenne T, Alexandre S. 2014. Characterization of membrane lipidome changes in *Pseudomonas aeruginosa* during biofilm growth on glass wool. PLoS ONE 9.

18. Aktas M, Danne L, Moller P, Narberhaus F. 2014. Membrane lipids in Agrobacterium tumefaciens: biosynthetic pathways and importance for pathogenesis. Front Plant Sci 5:109.

19. Wang S, Yu S, Zhang Z, Wei Q, Yan L, Ai G, Liu H, Ma LZ. 2014. Coordination of swarming motility, biosurfactant synthesis, and biofilm matrix exopolysaccharide production in Pseudomonas aeruginosa. Appl Environ Microbiol 80:6724–6732.

20. Ojha AK, Baughn AD, Sambandan D, Hsu T, Trivelli X, Guerardel Y, Alahari A, Kremer L, Jacobs WR, Jr., Hatfull GF. 2008. Growth of Mycobacterium tuberculosis biofilms containing free mycolic acids and harbouring drug-tolerant bacteria. Mol Microbiol 69:164–174.

21. Gianotti A, Serrazanetti D, Sado Kamdem S, Guerzoni ME. 2008. Involvement of cell fatty acid composition and lipid metabolism in adhesion mechanism of Listeria monocytogenes. Int J Food Microbiol 123:9–17.

22. Rowlett VW, Mallampalli V, Karlstaedt A, Dowhan W, Taegtmeyer H, Margolin W, Vitrac H. 2017. Impact of Membrane Phospholipid Alterations in Escherichia coli on Cellular Function and Bacterial Stress Adaptation. J Bacteriol 199.

23. Tan BK, Bogdanov M, Zhao J, Dowhan W, Raetz CRH, Guan Z. 2012. Discovery of a cardiolipin synthase utilizing phosphatidylethanolamine and phosphatidylglycerol as substrates. Proc Natl Acad Sci U S A 109:6.

24. Romantsov T, Culham DE, Caplan T, Garner J, Hodges RS, Wood JM. 2017. ProP-ProP and ProP-phospholipid interactions determine the subcellular distribution of osmosensing transporter ProP in Escherichia coli. Mol Microbiol 103:469–482.

25. Mileykovskaya E, Dowhan W. 2009. Cardiolipin membrane domains in prokaryotes and eukaryotes. Biochim Biophys Acta 1788:2084–2091.

26. Rajendram M, Zhang L, Reynolds BJ, Auer GK, Tuson HH, Ngo KV, Cox MM, Yethiraj A, Cui Q, Weibel DB. 2015. Anionic Phospholipids Stabilize RecA Filament Bundles in Escherichia coli. Mol Cell 60:374–384.

27. Maloney E, Madiraju SC, Rajagopalan M, Madiraju M. 2011. Localization of acidic phospholipid cardiolipin and DnaA in mycobacteria. Tuberculosis (Edinb) 91 Suppl 1:S150–155.

28. Majdalani N, Gottesman S. 2005. The Rcs phosphorelay: a complex signal transduction system. Annu Rev Microbiol 59:379–405.

29. Pannen D, Fabisch M, Gausling L, Schnetz K. 2016. Interaction of the RcsB Response Regulator with Auxiliary Transcription Regulators in Escherichia coli. J Biol Chem 291:2357–2370.

30. Majdalani N, Hernandez D, Gottesman S. 2002. Regulation and mode of action of the second small RNA activator of RpoS translation, RprA. Mol Microbiol 46:813–826.

31. Davalos-Garcia M, Conter A, Toesca I, Gutierrez C, Cam K. 2001. Regulation of osmC gene expression by the two-component system rcsB-rcsC in Escherichia coli. J Bacteriol 183:5870–5876.

32. Stout V, Gottesman S. 1990. RcsB and RcsC: a two-component regulator of capsule synthesis in *Escherichia coli*. J Bacteriol 172:11.

33. Lin TY, Weibel DB. 2016. Organization and function of anionic phospholipids in bacteria. Appl Microbiol Biotechnol 100:4255–4267.

34. Li C, Tan BK, Zhao J, Guan Z. 2016. In Vivo and in Vitro Synthesis of Phosphatidylglycerol by an Escherichia coli Cardiolipin Synthase. J Biol Chem 291:25144–25153.

35. Tropp BE, Ragolia L, Xia W, Dowhan W, Milkman R, Rudd KE, Ivanisevic R, Savic DJ. 1995. Identity of the *Eschericia coli cls* and *nov* genes. J Bacteriol 177:4.

36. Bury-Moné S, Nomane Y, Reymond N, Barbet R, Jacquet E, Imbeaud S, Jacq A, Bouloc P. 2009. Global analysis of extracytoplasmic stress signaling in Escherichia coli. PLoS Genet 5:e1000651.

37. Ferrieres L, Thompson A, Clarke DJ. 2009. Elevated levels of sigma S inhibit biofilm formation in Escherichia coli: a role for the Rcs phosphorelay. Microbiology 155:3544–3553.

38. Howery KE, Clemmer KM, Rather PN. 2016. The Rcs regulon in Proteus mirabilis: implications for motility, biofilm formation, and virulence. Curr Genet 62:775–789.

39. Clarke DJ. 2010. The Rcs phosphorelay: more than just a two-componen Future Microbiol 5:1173–1184.

40. Tomura A, Ishikawa T, Sagara Y, Miki T, Sekimizu K. 1993. Requirement of phosphatidylglycerol for flagellation of *Escherichia coli*. FEBS Lett 329:4.

41. Cho SH, Szewczyk J, Pesavento C, Zietek M, Banzhaf M, Roszczenko P, Asmar A, Laloux G, Hov AK, Leverrier P, Van der Henst C, Vertommen D, Typas A, Collet JF. 2014. Detecting envelope stress by monitoring beta-barrel assembly. Cell 159:1652–1664.

42. Konovalova A, Perlman DH, Cowles CE, Silhavy TJ. 2014. Transmembrane domain of surface-exposed outer membrane lipoprotein RcsF is threaded through the lumen of beta-barrel proteins. Proc Natl Acad Sci U S A 111:E4350–4358.

43. Dominguez-Bernal G, Pucciarelli MG, Ramos-Morales F, Garcia-Quintanilla M, Cano DA, Casadesus J, Garcia-del Portillo F. 2004. Repression of the RcsC-YojN-RcsB phosphorelay by the IgaA protein is a requisite for Salmonella virulence. Mol Microbiol 53:1437–1449.

44. Gold VAM, Robson A, Bao H, Romantsov T, Duong F, Collinson I. 2010. The action of cardiolipin on the bacterial translocon. Proc Natl Acad Sci U S A 107:6.

45. Schulze RJ, Komar J, Botte M, Allen WJ, Whitehouse S, Gold VA, Lycklama ANJA, Huard K, Berger I, Schaffitzel C, Collinson I. 2014. Membrane protein insertion and proton-motive-force-dependent secretion through the bacterial holo-translocon SecYEG-SecDF-YajC-YidC. Proc Natl Acad Sci U S A 111:4844–4849.

46. Renner LD, Weibel DB. 2012. MinD and MinE interact with anionic phospholipids and regulate division plane formation in Escherichia coli. J Biol Chem 287:38835–38844.

47. Sekimizu K, Kornberg A. 1988. Cardiolipin activation of dnaA protein, the initiation protein of replication in Escherichia coli. J Biol Chem 263:5.

48. Laganowsky A, Reading E, Allison TM, Ulmschneider MB, Degiacomi MT, Baldwin AJ, Robinson CV. 2014. Membrane proteins bind lipids selectively to modulate their structure and function. Nature 510:172–175.

49. Arias-Cartin R, Grimaldi S, Pommier J, Lanciano P, Schaefer C, Arnoux P, Giordano G, Guigliarelli B, Magalon A. 2011. Cardiolipin-based respiratory complex activation in bacteria. Proc Natl Acad Sci U S A 108:7781–7786.

50. Romantsov T, Guan Z, Wood JM. 2009. Cardiolipin and the osmotic stress responses of bacteria. Biochim Biophys Acta 1788:2092–2100.

51. Hiraoka S, Matsuzaki H, Shibuya I. 1993. Active increase in cardiolipin synthesis in the stationary growth phase and its physiological significance in Escherichia coli. FEBS Lett 336:4.

52. Mukhopadhyay R, Huang KC, Wingreen NS. 2008. Lipid localization in bacterial cells through curvature-mediated microphase separation. Biophys J 95:1034–1049.

53. Huang KC, Ramamurthi KS. 2010. Macromolecules that prefer their membranes curvy. Mol Microbiol 76:822–832.

54. van Elsas JD, Semenov AV, Costa R, Trevors JT. 2011. Survival of Escherichia coli in the environment: fundamental and public health aspects. ISME J 5:173–183.

55. Shrout JD, Chopp DL, Just CL, Hentzer M, Givskov M, Parsek MR. 2006. The impact of quorum sensing and swarming motility on Pseudomonas aeruginosa biofilm formation is nutritionally conditional. Mol Microbiol 62:1264–1277.

56. Capita R, Riesco-Pelaez F, Alonso-Hernando A, Alonso-Calleja C. 2014. Exposure of Escherichia coli ATCC 12806 to sublethal concentrations of food-grade biocides influences its ability to form biofilm, resistance to antimicrobials, and ultrastructure. Appl Environ Microbiol 80:1268–1280.

57. Wolfe AJ, Chang DE, Walker JD, Seitz-Partridge JE, Vidaurri MD, Lange CF, Pruss BM, Henk MC, Larkin JC, Conway T. 2003. Evidence that acetyl phosphate functions as a global signal during biofilm development. Mol Microbiol 48:977–988.

58. Castelli ME, Vescovi EG. 2011. The Rcs signal transduction pathway is triggered by enterobacterial common antigen structure alterations in Serratia marcescens. J Bacteriol 193:63–74.

59. Wang Q, Zhao Y, McClelland M, Harshey RM. 2007. The RcsCDB signaling system and swarming motility in Salmonella enterica serovar typhimurium: dual regulation of flagellar and SPI-2 virulence genes. J Bacteriol 189:8447–8457.

60. Schwan WR, Shibata S, Aizawa S, Wolfe AJ. 2007. The two-component response regulator RcsB regulates type 1 piliation in Escherichia coli. J Bacteriol 189:7159–7163.

61. Abraham SN, Babu JP, Giampapa CS, Hasty DL, Simpson WA, Beachey EH.1985. Protection against Escherichia coli-induced urinary tract infections with hybridoma antibodies directed against type 1 fimbriae or complementary D-mannose receptors. Infect Immun 48:625–628.

62. Hirakawa H, Nishino K, Hirata T, Yamaguchi A. 2003. Comprehensive studies of drug resistance mediated by overexpression of response regulators of two-component signal transduction systems in Escherichia coli. J Bacteriol 185:1851–1856.

63. Strahl H, Burmann F, Hamoen LW. 2014. The actin homologue MreB organizes the bacterial cell membrane. Nat Commun 5:3442.

64. Yang X, Sheng W, He Y, Cui J, Haidekker MA, Sun GY, Lee JC. 2010. Secretory phospholipase A2 type III enhances alpha-secretase-dependent amyloid precursor protein processing through alterations in membrane fluidity. J Lipid Res 51:957–966.

65. Finka A, Goloubinoff P. 2014. The CNGCb and CNGCd genes from Physcomitrella patens moss encode for thermosensory calcium channels responding to fluidity changes in the plasma membrane. Cell Stress Chaperones 19:83–90.

66. Unsay JD, Cosentino K, Subburaj Y, García-Sáez AJ. 2013. Cardiolipin effects on membrane structure and dynamics. Langmuir 29:10.

67. Nenninger A, Mastroianni G, Robson A, Lenn T, Xue Q, Leake MC, Mullineaux CW. 2014. Independent mobility of proteins and lipids in the plasma membrane of Escherichia coli. Mol Microbiol 92:1142–1153.

68. Tuson HH, Copeland MF, Carey S, Sacotte R, Weibel DB. 2013. Flagellum density regulates *Proteus mirabilis* swarmer cell motility in viscous environments. J Bacteriol 195:10.

69. Dubois-Brissonnet F, Trotier E, Briandet R. 2016. The Biofilm Lifestyle Involves an Increase in Bacterial Membrane Saturated Fatty Acids. Front Microbiol 7:1673.

70. McClelland M, Sanderson KE, Spieth J, Clifton SW, Latreille P, Courtney L, Porwollik S, Ali J, Dante M, Du F, Hou S, Layman D, Leonard S, Nguyen C, Scott K, Holmes A, Grewal N, Mulvaney E, Ryan E, Sun H, Florea L, Miller W, Stoneking T, Nhan M, Waterston R, Wilson RK. 2001. Complete genome sequence of Salmonella enterica serovar Typhimurium LT2. Nature 413:852–856.

71. Miller JH. 1972. Experiments in molecular genetics. Cold Spring Harbor Laboratory.

72. Datsenko KA, Wanner BL. 2000. One-step inactivation of chromosomal genes in Escherichia coli K-12 using PCR products. Proc Natl Acad Sci U S A 97:6640–6645.

73. Seidman CE, Struhl K, Sheen J, Jessen T. 2001. Introduction of plasmid DNA into cells. Curr Protoc Mol Biol Chapter 1:Unit1 8.

74. O’Toole GA, Kolter R. 1998. Initiation of biofilm formation in *Pseudomonas fluorescens* WCS365 proceeds via multiple, convergent signalling pathways: a genetic analysis. Mol Microbiol 28:13.

75. Belin D. 2010. In vivo analysis of protein translocation to the Escherichia coli periplasm. Methods Mol Biol 619:103–116.

76. Czolkoss S, Fritz C, Holzl G, Aktas M. 2016. Two Distinct Cardiolipin Synthases Operate in Agrobacterium tumefaciens. PLoS One 11:e0160373.

